# UNC-45A Weakens and Breaks MT Lattice Independent of its Effect on Non-Muscle Myosin II

**DOI:** 10.1101/2020.06.20.163048

**Authors:** Juri Habicht, Ashley Mooneyham, Asumi Hoshino, Mihir Shetty, Xiaonan Zhang, Edith Emmings, Qing Yang, Courtney Coombes, Melissa K. Gardner, Martina Bazzaro

## Abstract

In invertebrates, UNC-45 regulates myosin stability and functions. Vertebrates have two distinct isoforms of the protein: UNC-45B, expressed in muscle cells only and UNC-45A, expressed in all cells and implicated in regulating both Non-Muscle Myosin II (NMII)- and microtubule (MT)-associated functions. Here we show for the first time that: *a) in vitro* UNC-45A binds to the MT lattice and weakens its integrity leading to MT bending, breakage and depolymerization, *b)* in cells, UNC-45A overexpression causes loss of MT mass and increase in MT breakages, *c)* both *in vitro* and in cells, UNC-45A destabilizes MTs independent of its NMII C-terminal binding domain and destabilization occurs even in presence of the NMII inhibitor blebbistatin. These findings are consistent with a not mutually exclusive but rather dual role of UNC-45A in regulating NMII activity and MT stability.

Because many human diseases, from cancer to neurodegenerative diseases, are caused by or associated with deregulation of MT stability our findings have profound implications in both, the biology of MTs as well as the biology of human diseases and possible therapeutic implications for their treatment.

## Introduction

UNC-45, a highly conserved member of UCS (UNC-45/CRO1/She4p) family, was discovered in the early ‘70s when mutations in this gene caused *C. elegans* to have uncoordinated movements, hence the term “UNC” (Epstein and Thomson, 1974). It was later discovered that this phenotype was due to the role of UNC-45 as a myosin chaperone (Barral et al., 1998; Barral et al., 2002; Bernick et al., 2010; Etard et al., 2007; Etheridge et al., 2002; Geach and Zimmerman, 2010; Hutagalung et al., 2002; Lee et al., 2011). In the early 2000s it was shown that unlike lower organisms, vertebrates have two isoforms of UNC-45: UNC-45A and UNC-45B. The latter is located on chromosome 17 and is expressed in muscle cells only where, similarly to UNC-45, it is required for striated muscle myosin folding and assembly (Bujalowski et al., 2014). UNC-45A on the other hand, is located on chromosome 15, is expressed in all cells and has multiple functions.

We and others have shown that in mammalian cells, UNC-45A binds to and co-localizes with Non-Muscle Myosin II (NMII) (Bazzaro et al., 2007; Iizuka et al., 2015; Iizuka et al., 2017a) and the C-terminal NMII-binding domain of UNC-45A is required for myosin II folding, myosin II binding with actin (Guo et al., 2011; Ni et al., 2011), and stress fiber assembly (Lehtimaki et al., 2017). This includes studies from our laboratory showing that UNC-45A plays a role in regulating NMII-assisted functions including cytokinesis(Bazzaro et al., 2007), exocytosis in immune cells (Iizuka et al., 2015), axonal growth of neurons(Iizuka et al., 2017b) and tunneling nanotube formation(Lou et al., 2018).

In addition to its myosin-dependent functions, UNC-45A was recently found to colocalize with gamma-tubulin at the microtubule (MT) organizing center (MTOC), biochemically co-fractionates with gamma-tubulin (Jilani et al., 2015), and fractionates with well-known MT-associated and destabilizing proteins, including katanin and MCAK (Itzhak et al., 2016). Additionally, we have recently shown that UNC-45A is a novel Microtubule-Associated-Protein (MAP) with MT destabilizing properties, and that UNC-45A is overexpressed in paclitaxel-resistant human cancer cells where it counteracts the MT stabilizing effects of paclitaxel (Habicht et al., 2019b; Mooneyham et al., 2019). While UNC-45A has been most commonly associated with actomyosin related function, recent studies contribute to the growing body of evidence that UNC-45A has myosin-independent functions and regulates MTs stability.

A number of MAPs are involved in regulating MT stability via either promoting polymerization or depolymerization from MT ends (Goodson and Jonasson, 2018). A special class of MT destabilizing proteins, known as MT severing proteins, exert destabilization from the microtubule lattice. MT severing proteins bind along the length of MTs, causing morphological changes to the MT lattice which is followed by MT breakage into shorter fragments which will eventually depolymerize (Hartman et al., 1998; Hartman and Vale, 1999).

In this study we show for the first time that UNC-45A depolymerizes paclitaxel-stabilized MTs by binding to their lattice and weakening it which causes kinks and breakages along the MTs length. This is followed by MT depolymerization in a dose dependent manner. We also show for the first time that both *in vitro* and in cells, the C-terminal NMII-binding domain of UNC-45A is not required for MT binding nor for their destabilization. Lastly, we show that in cells, UNC-45A depolymerizes MTs independent of its effect on NMII since UNC-45A-mediated MT depolymerization occurs even in presence of the NMII inhibitor blebbistatin. Taken together our studies support the role of UNC-45A as a novel member of the MT destabilizing protein via weakening, bending and breaking MT lattice, and its dual and not mutually exclusive role in regulating actomyosin and MT activities.

## Materials and Methods

### Preparation of paclitaxel-stabilized, rhodamine-labeled microtubules

The paclitaxel-stabilized rhodamine-labeled MTs were prepared as we have previously described (Mooneyham et al., 2019). Briefly, unlabeled tubulin (cytoskeleton #T240) and rhodamine labeled tubulin (cytoskeleton #TL590M) were mixed in a 5:1 ratio in Brb80 buffer with 1mM DTT and 1mM GTP then incubated on ice for 5 minutes. Next, the mixture was incubated at 37^°^C where 1/10 of the reaction mixture volume of 1μM, 10μM, and 100μM paclitaxel (Cytoskeleton #TXD01) were added sequentially for 5 minutes each to create labeled, taxol-stabilized microtubules at a final concentration of 40μM. After incubation, the MT mixture was diluted with warm Brb80 solution containing 100μM paclitaxel and 1mM DTT to be flowed into the TIRF chamber. All microtubules were prepared on the same day of the experiment.

### Construction and preparation of flow chambers for TIRF imaging

A 22mm × 50mm glass coverslip was used as the base of the chamber and an 18mm × 18mm coverslip was used as the top. Prior to chamber assembly, coverslips were thoroughly cleaned and silanized. Three narrow strips of parafilm were stacked in three equidistant columns on the base coverslip to create two separate experimental chambers per slide. Once assembled, vacuum-assistance was used to flow an anti-rhodamine antibody solution in Brb80 into the chambers. After 10 minutes of room temperature incubation, the antibody solution was flushed out with two chamber volumes of Brb80 and a blocking solution of 1% PF127 in Brb80 was introduced. After 10 minutes of incubation the chambers were flushed with 2 channel volumes of Brb80 and the rhodamine-labeled MT mixture was added and allowed to attach to the antibody-coated coverslip for 15 minutes.

### UNC-45A-GFP microtubule binding and impact on MT kinking and MT mass

For UNC-45A-GFP MT binding, a final reaction mixture, containing 1x imaging buffer and 0.6 μM final concentration of UNC-45A-GFP, was introduced into the imaging chamber and the interaction between UNC-45A-GFP and Paclitaxel-stabilized microtubule was visualized via 488 nm and 561 nm lasers generated from Nikon™ TI-TIRF-PAU illuminator, which provided Total Internal Reflection Fluorescence (TIRF) illumination. The images were collected from a Nikon™ CFI Apo TIRF 100x oil objective using an Andor™ iXon EMCCD camera. For the impact of UNC-45A-GFP on MT kinking and MT mass, a final reaction mixture, containing imaging buffer and 125nM or 250nM final concentration of UNC-45A-GFP, was introduced into the TIRF imaging chamber, and the interaction between UNC-45A-GFP and paclitaxel-stabilized microtubules was visualized via time-lapse imaging 488 nm and 561 nm lasers generated from the Zeiss TIRF microscope at 100X magnification. Alternatively, images were taken after 10 minute and 20 minutes incubation with UNC-45A-GFP in each condition. A kinked microtubule was defined as an angle observed within an individual microtubule. Microtubule mass was measured as average microtubule fluorescent intensity from 3 individual areas per field of view. Laser power and exposure time were minimized while TIRF angle was maximized to avoid photobleaching and photodamage.

### Recombinant protein

GFP-tagged UNC-45A full-length (UNC-45A WT, 1-944 aa), its C-terminal (dC-UNC-45A,1- 554 aa) or N-terminal (dN-UNC-45A, 125-944 aa) were cloned into pGEX-2TK to generate the GST-GFP-UNC-45A protein. The protein was expressed in Rosetta (DE3) pLysS and following GST removal it was affinity purified and dialyzed as we have previously described (Mooneyham et al., 2019).

### *In vitro* tubulin polymerization assay

The tubulin polymerization assay was performed as instructed in the Cytoskeleton Tubulin Polymerization Assay Kit (#BK006P) with the following modifications: no glycerol was used as a polymerization enhancer due to its known interference with tubulin ligand binding and instead, a 5mg/mL starting concentration of purified tubulin was used to enhance spontaneous tubulin polymerization. Assay was performed with 25nM or 75nM of purified UNC-45A protein (wildtype, C-terminally deleted, or N-terminally deleted) and 1μM or of nocodazole as a depolymerization control. Each polymerization curve was analyzed as a percentage of the control polymerization curve. All experiments performed in triplicate using at least two different batches of recombinant protein (specifically, we used 3 different deltaN batches and 2 different batches of WT and deltaC).

### Cell culture

Original and UNC-45A knockout (KO) human osteosarcoma U2OS cells were a generous gift of Drs. Drs. Jaakko Lehtimäki and Pekka Lappalainen and were cultured as previously described (Lehtimaki et al., 2017). Rat RFL-6 fibroblasts were purchased from the American Type Culture Collection (ATCC) and cultured in Ham’s F-12K medium (Thermo Fisher) supplemented with 20% FBS as previously described (Qiang et al., 2006). All cells were recently authenticated and routinely tested negative for mycoplasma.

### Antibodies and chemicals

Rabbit anti UNC-45A (Protein Tech, 1956-1-AP) raised against the C-terminus of the human UNC-45A (Protein Tech, 1956-1-AP) was used for Western blot and immunofluorescence as we have previously described (Habicht et al., 2019a; Mooneyham et al., 2019). Mouse polyclonal anti UNC-45A raised against full length UNC-45A (Abnova, H00055898-B01P) was used for Western blot to detect recombinant UNC-45A and its mutants (Figure 3B). Mouse anti-α-tubulin (Sigma-Aldrich, T6074), for Western blot, rabbit polyclonal α-tubulin (Abcam, ab18251), mouse monoclonal anti-detyrosinated-α-tubulin (EMD Millipore AB3201), rabbit monoclonal anti-GFP (Thermo Fisher Scientific G10362), mouse monoclonal anti-flag (Sigma-Aldrich, F1804), Peroxidase-linked anti-mouse immunoglobulin G and peroxidase-linked antirabbit immunoglobulin G (both Cytiva, formerly known as GE Healthcare Bio-Sciences NA931 and NA934), Alexa Fluor 594-conjugated Donkey Anti-Mouse IgG (1:250) and FITC-conjugated Goat Anti-Rabbit IgG (1:200; both Jackson ImmunoResearch Laboratories 715-585-150 and 111-095-003). Paclitaxel was purchased from Teva Pharmaceuticals. Blebbistatin was purchased from Sigma-Aldrich, St. Louis, MO. Tubulin Tracker Deep Red was purchased from Thermo Fisher Scientific (Waltham, MA) cat # T34077.

### Modulation of UNC-45A and katanin expression levels in cells

For UNC-45A silencing and overexpression scramble and UNC-45A shRNAs lentiviral supernatant or empty vector control and UNC-45A-GFP lentiviral supernatants were prepared and used to infect RFL-6 cells as we have previously described (Iizuka et al., 2015; Iizuka et al., 2017a; Mooneyham et al., 2019). For ectopic expression, RFL-6 or U2OS cells were infected with lentiviral particle carrying FLAG only (control) or FLAG-tagged UNC-45A WT, deltaN and deltaC and expression levels were evaluated 24- or 48- hours post infection. For ectopic expression of UNC-45A in RFL-6 cells, cells were infected with either GFP or GFP-UNC-45A lentiviral supernatants as we have previously described (Mooneyham et al., 2019). For ectopic expression of katanin, RFL-6 cells were transduced with GFP or GFP-katanin plasmids using Fugene HD.

### Western blot analysis

Total cellular protein (10–50 μg) from each sample was separated by SDS-PAGE, transferred to PVDF membranes and subjected to Western blot analysis using the specified antibodies. Amido black staining was performed to confirm equal protein loading.

### Blebbistatin treatment

RFL-6 cells were treated with either 12.5 or 25 μM blebbistatin or DMSO for 1 hour before lysates were made using RIPA buffer (50 mM Tris 7.5, 150 mM NaCl, 1% NP-40, 0.5% sodium deoxycholate, 0.1% SDS) as previously described (Joo and Yamada, 2014).

### Immunofluorescence microscopy, image acquisition, and analysis

For co-localization analysis of UNC-45A and its mutants and MTs, U2OS cells were fixed in cold methanol for 5 minutes at −20°C. After blocking with 5% BSA in PBST, cells were stained with anti-UNC-45A and anti-α-tubulin primary antibodies followed by FITC- or Alexa Fluor 594-conjugated secondary antibodies and analyzed via confocal fluorescence microscopy. Images were taken with an Olympus BX2 upright microscope. A UPlanApo N 60X/1.42 NA objective was used. FITC was excited with a 488 nm laser and emission collected between 505 and 525 nm. For Alexa Fluor a 594 nm laser was used for excitation and emission collected between 560 and 660 nm. Images were taken with sequential excitation. Co-localization analysis was done using the Fiji 2 software Coloc 2 plugin and Pearson’s correlation coefficient (PCC) and calculated as previously described (Dunn et al., 2011). For studies on MTs morphology (breakages) and mass following UNC-45A overexpression, GFP-control of GFP-UNC-45A expressing RFL-6 cells were extracted in a microtubule stabilizing buffer with 0.5% Triton-X for 30 seconds to remove free tubulin and then fixed with 0.5% glutaraldehyde for 10 minutes. After fixation, cells were quenched for 7 minutes with 0.1% NaBH4 and then rehydrated in PBS before blocking in AbDil (2% TBST with 2%BSA and 0.1% Azide) for 30 minutes as previously described (Ritter et al., 2017). After blocking, cultures were stained with anti-α- tubulin and anti-GFP primary antibody followed by Alexa Fluor 594 and FITC secondary antibodies. Cells were then rinsed in TBST and mounted in a medium that reduces photobleaching. Images were obtained on an Axiovert 200 microscope (Zeiss, Thornwood, NY) equipped with a high-resolution CCD camera. All images were obtained using identical camera, microscope, and imaging criteria such as gain, brightness, contrast, and exposure time. Digital gray values of image pixels representing arbitrary fluorescence units (AFUs) were obtained using FiJi software. For microtubule mass measurement in RFL-6 cells, quantification was performed on three comparable, equally sized areas per cell. For the analysis of microtubule breakages microtubule free ends were counted per cell as previously described (Ahmad et al., 1999).

### Live imaging of MTs in RFL-6 cells

To determine the effects of UNC-45A overexpression on MTs bending and breaking in live cells, GFP-control of GFP-UNC-45A overexpressing RFL-6 cells were treated with Tubulin Tracker Deep Red according to the manufacture’s recommendation. Tubulin Tracker Deep was diluted 1:2000. This dilution corresponds to 500 nM of taxol (personal communication from the Thermo

Fisher technical support). Cells (GFP and GFP-UNC-45A) were treated with Tubulin Tracker Deep Red sequentially so the imaging was done under the same conditions. Timelapse fluorescent images were collected with a Zeiss Axio observer Z1 inverted microscope using a 100X/1.46 for objective lens. Digital images were collected every 8 seconds intervals over the period of 8 minutes using a Cy5 filter cube. Laser power and exposure time were minimized to avoid photobleaching and photodamage and all images (for GFP and GFP-UNC-45A) were taken under the same conditions. A bending MT was defined as a MT in which bending was observable throughout the time of imaging.

## Data availability

### Statistical analysis

Results are reported as mean ± Standard Deviation of three or more independent experiments. Unless otherwise indicated, statistical significance of difference was assessed by two-tailed Student’s t using Prism (V.4 Graphpad, San Diego, CA) and Excel. The level of significance was set at p < 0.05.

## Results

### *In vitro*, UNC-45A-mediated MT destabilization is preceded by kinks and breakages in the MT body

We have recently shown that UNC-45A is a novel Microtubule-Associated-Protein (MAP) that binds and destabilizes MTs *in vitro* and *in vivo* in cancer cells and in fibroblasts (Habicht et al., 2019b; Mooneyham et al., 2019). To gain insights into the mechanism through which UNC-45A regulates MT stability, we used TIRF microscopy to examine the localization of UNC-45A on paclitaxel-stabilized rhodamine-labeled MTs of varying lengths. By sorting the microtubules by length and then quantifying the average localization of UNC-45A-GFP on the microtubules, we found that UNC-45A-GFP binds robustly along the length of the *in-vitro* microtubules, with a slight increase in UNC45A-GFP binding at microtubule ends (Figure 1A and B). This suggests that *in vitro*, UNC-45A may not act like a plus end MT destabilizing protein. While performing these experiments, we noticed that not only did MTs shorten in presence of UNC-45A in dose- and time-dependent fashion (Mooneyham et al., 2019), but they also appeared to be kinked and have gaps that were generally nonapparent in MTs in absence of UNC-45A (Figure 1C, arrows). Because this suggests that UNC-45A may destabilize the microtubule lattice, we next examined the events that preceded MT depolymerization in presence of UNC-45A. For this, paclitaxel-stabilized, rhodamine-labeled bovine-brain microtubules were incubated in absence and in presence of 125 or 250 nM of UNC-45A-GFP and the presence of kinks and overall MTs mass was recorded at 10 and 20 minutes of incubation. We found that the UNC-45A causes dose- and time-dependent increase in the numbers of MTs with kinks (Figure 2A) and a decrease in MT mass as compared to control condition (absence of UNC-45A). Timelapse TIRF microscopy of paclitaxel-stabilized, rhodamine-labeled bovine-brain MTs in presence of 250 nM of GFP-UNC-45A also shows that in presence (but not in absence) of UNC-45A MT breakages and depolymerization is preceded by MT kinks (Figure 2B). Quantification of number of kinked MTs and MT mass in absence and in presence of increasing concentrations of UNC-45A over time is given in Figure 2C and D respectively. Taken together this suggests that UNC-45A binds, kinks and destabilizes paclitaxel-stabilized MTs in a manner that is consistent with destabilizing MTs along their length.

**Figure 1.**
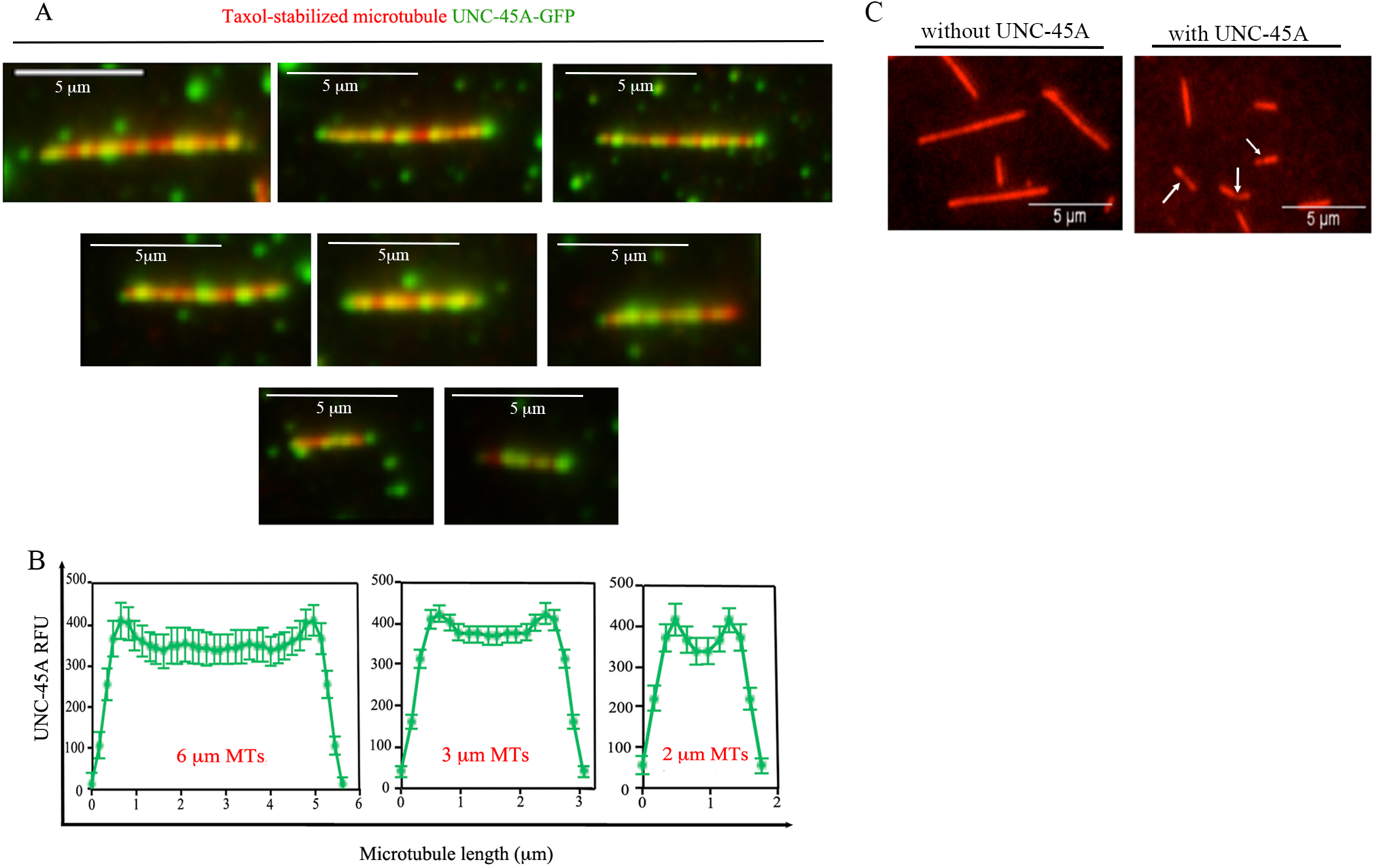
UNC-45A binds to MTs independent of their length. A. Sample images of paclitaxel-stabilized MTs (red) and UNC-45A-GFP (green) for different MT lengths. B. Quantification of UNC-45A-GFP localization on MTs, for three different average MT lengths (errors bars, SEM). C. D. Sample image of MTs with kinks (indicated by white arrows) in presence of 250 nM UNC-45A.

**Figure 2.**
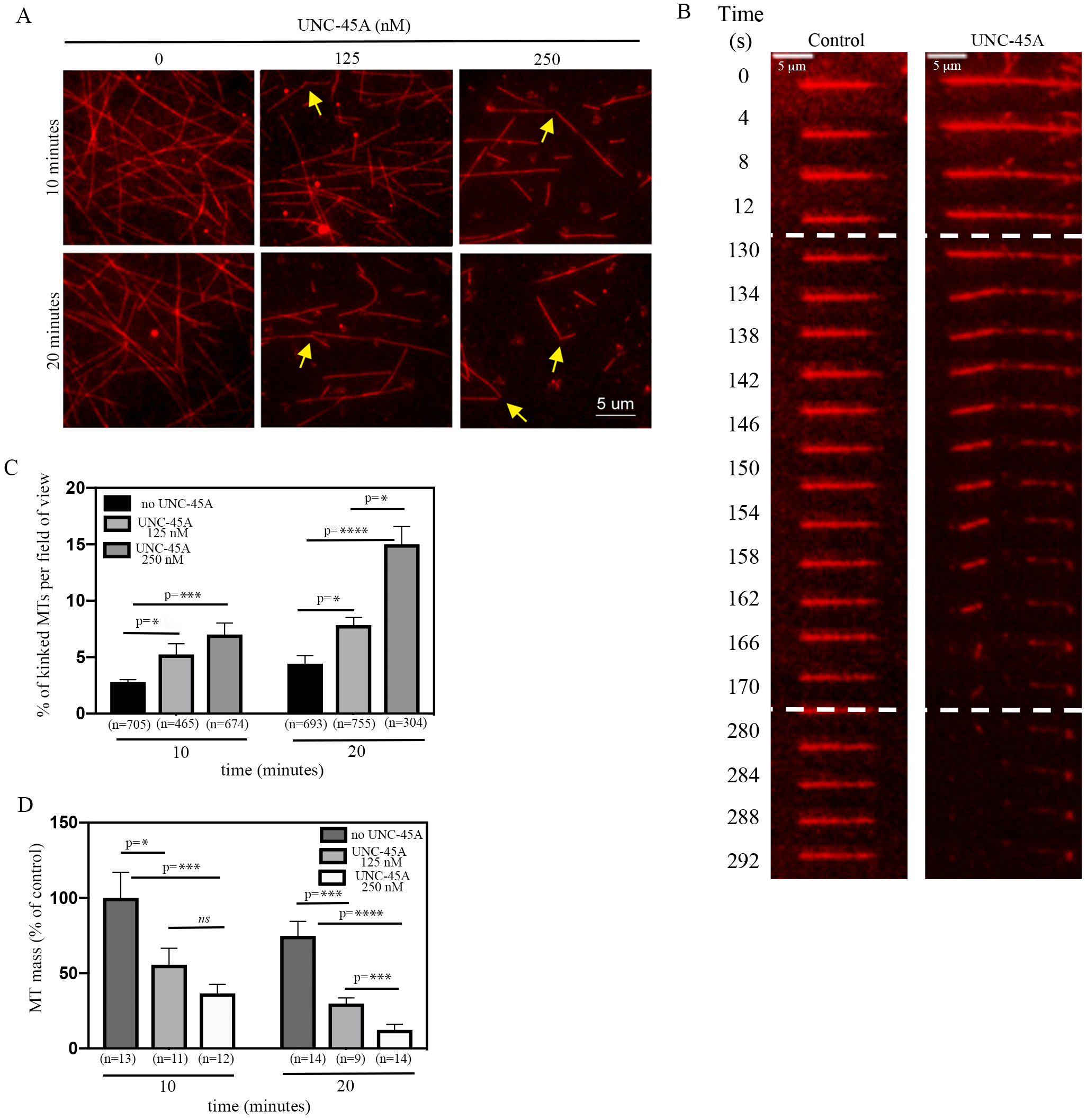
UNC-45A binds, weakens, kinks and destabilizes MTs *in vitro* in a concentration dependent manner. A. Sample images of paclitaxel-stabilized MTs in absence and in presence of 125 or 250 nM of UNC-45A-GFP for 10 and 20 minutes. Yellow arrows indicate MTs with kinks. B. Sample images of timelapse TIRF microscopy of paclitaxel-stabilized MTs in presence or in absence (control) of 250 nM of UNC-45A-GFP. Dotted lines indicate the timeframe in which MTs kink and break. C. MTs with kinks evaluated in absence and in presence of increasing concentrations of UNC-45A at 10 and 20 minutes of incubation. Results are expressed as % of kinked MTs per field of view (n=MTs evaluated per condition). D. MT mass in absence and in presence of increasing concentrations of UNC-45A at 10 and 20 minutes of incubation and determined by measuring the average microtubule fluorescent intensity from 3 individual areas per field of view (n= field of view measured per each condition). Results are expressed as a % of control.

### UNC-45A weakens and breaks paclitaxel-stabilized MTs *in vitro* independent of its C-terminal NMII-binding domain

Structurally, UNC-45A can be divided in four domains: an N-terminal domain, which contains three tetratricopeptide repeat (TPR) sequences and has been shown to bind to Hsp90; a central domain of largely unknown function; a neck domain, which has been recently proposed to be required for UNC-45 (worm isoform of UNC-45A) oligomerization; and a C-terminal, UCS domain that is critical for interaction with NMII (Barral et al., 1998; Ni et al., 2011; Shi and Blobel, 2010) (Figure 3A). We have shown that UNC-45A destabilizes paclitaxel-stabilized MTs in absence in any additional cellular components, including NMII, suggesting that *in vitro* UNC-45A destabilizes MTs independent of the presence of NMII(Mooneyham et al., 2019). Here we set out to determine which domain of UNC-45A is required for binding and destabilization of paclitaxel-stabilized MTs *in vitro*. We started by evaluating the MT binding abilities of recombinant UNC-45A WT, deltaN- or deltaC UNC-45A (Figure 3B) via TIRF microscopy. As we had previously shown (Mooneyham et al., 2019), UNC-45A is capable of directly binding MTs in a purified *in vitro* system without any additional cellular factors. The binding pattern is punctate and visible along the entire length of the MTs (Figure 3C, *top panel).* Similarly, and despite being approximatively half the size of the UNC-45A WT, the deltaC UNC-45A retained its ability to bind MTs efficiently and with a similar punctate pattern (Figure 3C, *bottom panel).*Conversely, deletion of the N-terminus domain completely abrogated the MT binding ability of UNC-45A (Figure 3C, *middle panel*). This suggests that the NMII-interacting domain is not required for UNC-45A’s MT-associated function and that the UNC-45A MT-interacting domain is located in the first 124 aa at the amino terminus. To correlate these findings with MT destabilizing activity, we determined the ability of recombinant full length (WT) UNC-45A, deltaN-or deltaC UNC-45A to limit net polymerization in a standard turbidimetric assay (Shelanski, 1973; Shelanski et al., 1973). We found that both, WT- and deltaC-UNC-45A caused a dose-dependent inhibition of net MT assembly *in vitro* (Figure 3D, *first and third panel*) while the deltaN-UNC-45A did not (Figure 3D, *second panel).* Importantly, UNC-45A acts on MTs at concentrations similar to the ones reported for other MT depolymerizing proteins when used in this assay (Di Paolo et al., 1997; Ng et al., 2006). Nocodazole was used as a positive control (Figure 3D, *last panel).* Furthermore, both full length and dC-UNC-45A do not seem to interfere with either the nucleation or the elongation phase, but rather with the MT steady state levels suggesting that UNC-45A acts to destabilize polymerized MTs. Importantly, because in this assay MTs are not stabilized by paclitaxel, this also suggests that UNC-45A does not exclusively depolymerize MTs by disrupting paclitaxel stabilization. Taken together, these results support the idea that UNC-45A has multiple distinct roles mediated by distinct protein domains.

**Figure 3.**
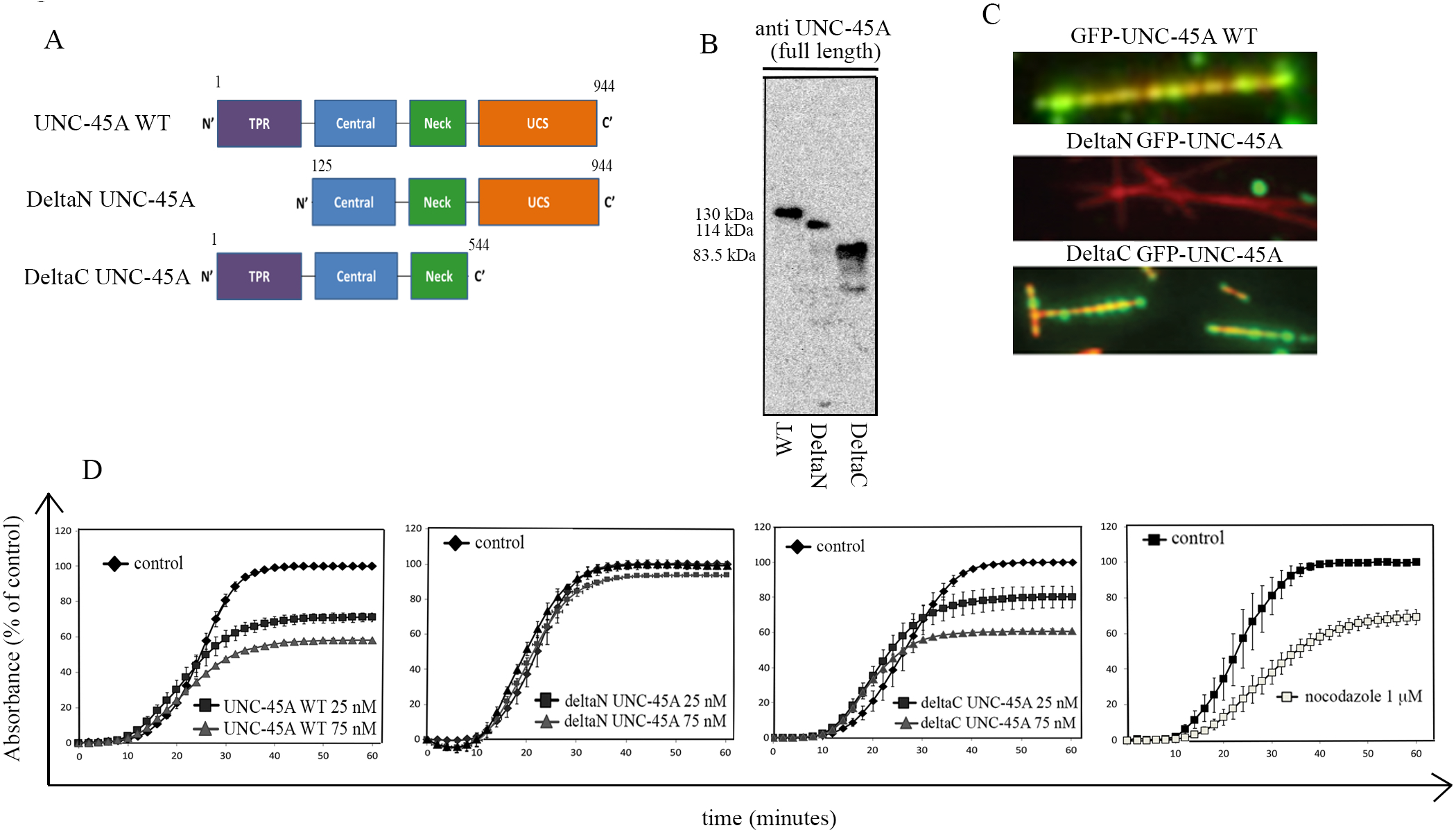
UNC-45A weakens and breaks MTs *in vitro* independent of its C-terminal myosin II-binding domain. A. Schematic diagram of full length (WT) UNC-45A (aa 1-944), deltaN UNC-45A (aa 125-944) or deltaC UNC-45A (aa 1-544). B. Recombinant full length (WT) UNC-45A-GFP (MW 130 kDa), deltaN UNC-45A-GFP (MW=114 kDa) and deltaC UNC-45A-GFP (83.5 kDa) were separated on a 4-15% SDS gel, transferred onto PVDF membrane and blotted against a rabbit polyclonal antibody raised against full length human UNC-45A. C. Representative images of WT, (*top*), C-terminally deleted (*middle*) or N-terminally deleted (*bottom*) UNC-45A (GFP-tagged) and rhodamine-labeled paclitaxel stabilized MTs obtained via TIRF microscopy. D. Tubulin 5mg/ml was incubated in absence (vehicle) or in presence of fulllength (WT) (*first panel*), deltaN (*second panel*) or deltaC-UNC-45A (*third panel*) or nocodazole (*panel panel*) at the indicated concentrations at 4°C. MT polymerization was induced at 37°C and optical density was recorded over a period of 1 hour. Results are expressed as optical density % of control. All experiments were performed in triplicates.

### The C-terminal NMII-binding domain of UNC-45A is not required for MT binding in cells

Having shown that in a cellular free system the N-terminal domain of UNC-45A is required for MT binding, we next asked whether the same is true in the cellular system. In order to test whether the N-terminal domain on UNC-45A is required for localization with MTs in cells we took advantage of the previously characterized UNC-45A knock-out (KO) U2OS cells (Lehtimaki et al., 2017). This is for two main reasons: *a*) because given the relatively high expression levels of UNC-45A in cells, ectopic expression of dominant negatives in UNC-45A WT cells is likely to result in high background and *b*) because ectopic expression of MAPs in general is known to cause “off-target” subcellular localization (Lehtimaki et al., 2017). First, we confirmed UNC-45A KO in U2OS cells via Western blot analysis. As shown in Figure 4A, in both the clones used (clone #5 and clone #6) (Lehtimaki et al., 2017) UNC-45A protein is not detected. We then ectopically expressed UNC-45A WT and its deltaN- and deltaC-mutants as FLAG-tag proteins in UNC-45A KO U2OS cells clone #6 and found that both UNC-45A WT and its mutants were expressed at similar levels in cells (Figure 4B). Next, we determined the subcellular localization of endogenously expressed UNC-45A (U2OS original) and ectopically expressed UNC-45A WT and its mutants with respect to MTs via IF microscopy as we have previously done (Habicht et al., 2019b; Mooneyham et al., 2019). As show in Figure 4C (*first panel from the top*) we found that, similarly to what we have previously described in other cell types, UNC-45A co-localizes with MTs in U2OS cells. Of note, we found that in this particular cell type, the MT network is particularly dense in the perinuclear region. We also found that similarly to endogenous UNC-45A, ectopically expressed FLAG-tagged UNC-45A and deltaC-UNC-45A colocalizes with MTs in U2OS cells (*Figure 4C, second and forth panels respectively).* Deletion of the N-terminal on the other hand resulted with a significant decrease in the levels of co-localization with MTs (*Figure 4C, third panel).* Quantification of co-localization between UNC-45A and its mutants and MTs is given in Figure 4D. Thus, even if at a lesser extent than in the cellular free system (*Figure 3, middle panel*), the N-terminus of UNC-45A seems to be important for UNC-45A localization to MTs in cells, strengthening our results that the UNC-45A’s NMII-binding domain is not required for MT binding.

**Figure 4.**
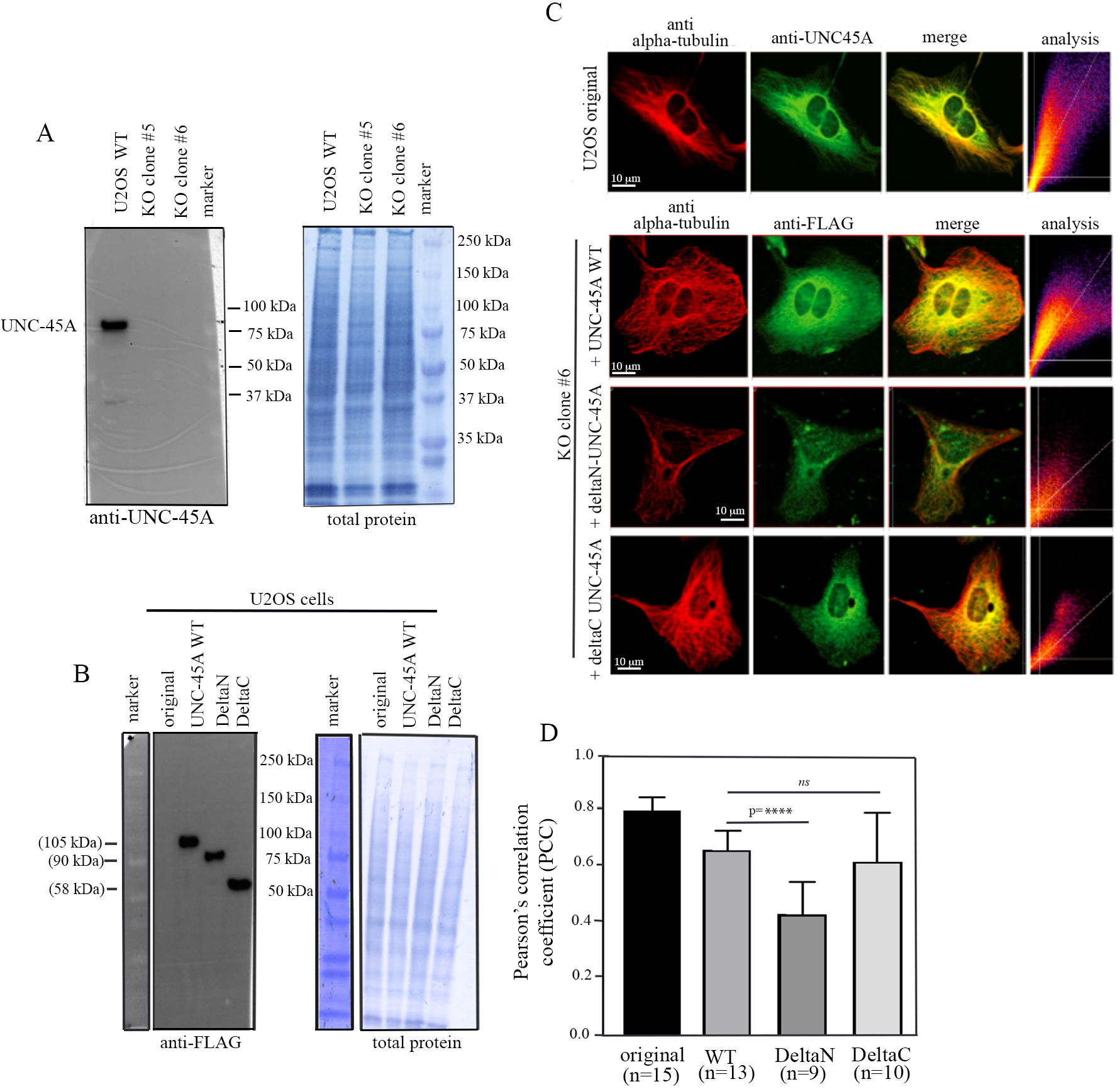
UNC-45A and its C-deleted but not N-deleted mutant co-localize with MTs in cells. A. *Left*, Western blot analysis for levels of UNC-45A in original and UNC-45A knock-out (KO) U2OS cells using anti-UNC-45A antibody. Two different clones were used. *Right*, total protein (amido black staining) was used as loading control. B. *Left*, Western blot analysis for levels of UNC-45A in original and UNC-45A KO clone #6 ectopically expressing either UNC-45A WT or deltaN-UNC-45A or dentaC-UNC-45A as a FLAG-tagged proteins using an anti-FLAG antibody. *Right*, Total protein (amido black staining) was used as loading control. C. *First panels from the top.* Representative images of tubulin (red) and UNC-45A (green) in original U2Os cells. Yellow is the merged image. Co-localization is expressed as PCC. *Second panels from the top.* Representative images of tubulin (red) and UNC-45A WT (FLAG-tagged) ectopically expressed in UNC-45A KO U2Os (clone #6). Yellow is the merged image. Co-localization is expressed as PCC. *Third panels from the top.* Representative images of tubulin (red) and deltaN UNC-45A (FLAG-tagged) ectopically expressed in UNC-45A KO U2Os (clone #6). Yellow is the merged image. Co-localization is expressed as PCC. *Bottom panels*. Representative images of tubulin (red) and deltaC UNC-45A (FLAG-tagged) ectopically expressed in UNC-45A KO U2Os (clone #6). Yellow is the merged image. Co-localization is expressed as PCC. D. Quantification of co-localization between UNC-45A and its mutants and MTs as determined via Pearson’s correlation coefficient (PCC). n= number of cells evaluated per each condition (3 different area per cell were evaluated).

### UNC-45A overexpression in RFL-6 cells leads to loss of MT mass and increase in MT breakages

We first determined whether, similarly to what we have shown for other cell types (Habicht et al., 2019a; Mooneyham et al., 2019), UNC-45A destabilizes MT in the rat lung fibroblasts RLF-6 cells. We used these cells because they have a very distinct MT network that facilitates analysis of MTs and their mass (Baas and Sudo, 2010; Sudo and Baas, 2010). As shown in Figure 5A (*left panel*) overexpression of UNC-45A led to an approx. 50% decrease in the levels of detyrosinated alpha tubulin, a marker of long-lived microtubules, in lysates of RFL-6 cells. Complementary experiments showed no differences in the levels of detyrosinated tubulin in UNC-45A KD cells versus control (Figure 5A, *right panel*). This could be due to the fact that RFL-6 has a relative stable morphology as compared to other cell types and that there is a direct correlation between stable morphology and MTs stability (Baas et al., 2016; Qiang et al., 2006). Next, we determined the effects of UNC-45A overexpression in RFL-6 via IF by quantifying the MT mass in control versus overexpressing cells. Specifically, following GFP or UNC-45A-GFP overexpression, cells were fixed, extracted and stained with anti-alpha tubulin antibody 24 hours post infection. Extraction was done as previously described to minimize the background due to free tubulin within cells (Sudo and Baas, 2011). As shown in Figure 5B, we found that ectopic expression of UNC-45A results in a significant decrease in total MT mass as compared to control cells. Quantification of MT mass per each condition is given in Figure 5C. Interestingly, we found no difference in MT mass in “low” versus “high” UNC-45A expressing RFL-6 cells (Supplementary Figure 1A and B). The difference in GFP expression levels in GFP-UNC45A “low” and GFP-UNC-45A “high” expressing RFL-6 was consistently >60%. Together this suggests that even a modest increase in UNC-45A is sufficient to cause a saturating effect. The loss of MT mass following UNC-45A overexpression was similar to the one that we observed by overexpressing the known MT severing protein katanin (Qiang et al., 2006) in the same cell type (Figure 5D and E). Importantly, we found that, also similarly to what has been previously reported for katanin (Qiang et al., 2006), UNC-45A overexpression resulted in MT breakages at the edges of RLF-6 cells as compared to control (Figure 5F and G, and Supplementary Figure 1C). Lastly, we confirmed that UNC-45A overexpression leads to MT breakages in live cells. Timelapse microscopy of Deep Red Tubulin Tracker-labelled MTs shows that GFP-UNC-45A overexpression in RFL-6 cells results with increased MT bending and breakages as compared to GFP transfected control cells (Figures 5H and I). Quantification of number of MT breakages per cell (three fields of view per cells were counted) in each condition is given in Figure 5J. We found that not all MT breakages were preceded by visible MT bending but found that the number of MT breakages that were preceded by bending is significantly higher in GFP-UNC-45A overexpressing cells as compared to control. Quantification of MTs bending and breakages per each condition was done in cells expressing similar amount of GFP (shown in Figure 5L). Importantly, the experiments in live cells were conducted in presence of 500 nM of taxol (personal communication from the manufacturer) confirming our earlier results that in cells, UNC-45A counteracts the MT stabilizing effect of paclitaxel (Mooneyham et al., 2018).

**Figure 5.**
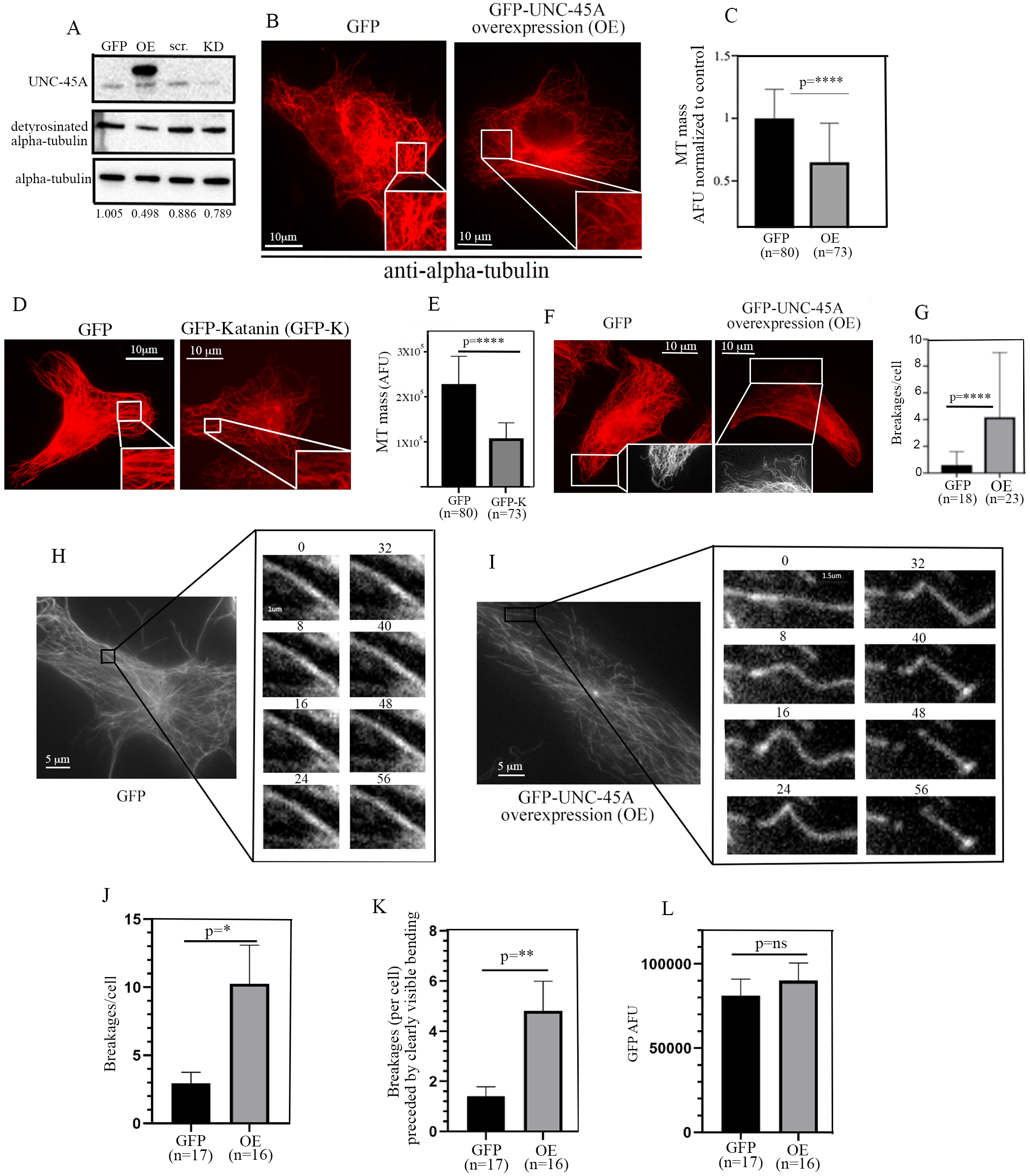
UNC-45A overexpression leads to loss of MT mass and increase in MT breakages in RFL-6 cells. A. Western blot analysis for levels of UNC-45A and detyrosinated alpha-tubulin (here used as a marker of MT stability) in GFP-UNC-45A overexpressing (OE) or UNC-45A knockdown (KD) RFL-6 cells as compared to their relative controls GFP and scramble (scr.). Total alpha-tubulin was used as a loading control. Numbers represent the ratio between detyrosinated alpha-tubulin and total alpha-tubulin per each condition. B. Representative images of GFP and GFP-UNC-45A expressing RFL-6 cells fixed, and stained with an anti-alpha-tubulin antibody to visualize MT mass. All images were taking using the same exposure time in cells expressing the similar amounts of GFP. C. Quantification of MT mass per each condition expressed as Arbitrary Fluorescence Units (AUF). n=number of cells analyzed per each condition (3 different areas per cell were evaluated). D. Representative images of GFP and GFP-Katanin (GFP-K) expressing RFL-6 cells fixed, and stained with an anti-alpha-tubulin antibody to visualize MT mass. All images were taken using the same exposure time in cells expressing similar amounts of GFP. E. Quantification of MT mass per each condition expressed as Arbitrary Fluorescence Units (AUF). n=number of cells analyzed per each condition (3 different areas per cell were evaluated). F. Representative images of GFP and GFP-UNC-45A expressing RFL-6 cells fixed, and stained with an anti-alpha-tubulin antibody to visualize MTs and their breakages. All images were taken using the same exposure time in cells expressing similar amounts of GFP. Inset are black and white details. G. Quantification of MT breakages per cell per each condition. n=number of cells analyzed per each condition. H. Live cell sample images of MTs (shown in white) in GFP transfected RFL-6 cells. Eight sequential timelapse frames are shown taken 8 seconds apart. I. Live cell sample images of MTs (shown in white) in GFP-UNC-45A overexpressing (OE) RFL-6 cells. Eight sequential timelapse frames are shown taken 8 seconds apart. J. Quantification of MT breakages per cell per each condition. K. Quantification of MT breakages per cell that were preceded by clearly visible bending per each condition. n= number of cells analyzed. L. Quantification of GFP (green signal) intensity in GFP and GFP-UNC-45A cells expressed as Arbitrary Fluorescence Units (AFU).

### In RFL-6 cells, UNC-45A weakens and breaks MT lattice in absence of its C-terminal myosin II binding domain, including in the presence of the myosin II inhibitor blebbistatin

Having established that in RFL-6 cells, UNC-45A destabilizes MTs via weakening and bending of MT lattice, we next asked whether the C-terminal NMII-binding domain of UNC-45A is required for this effect. While performing these experiments we noticed that although UNC-45A WT is overexpressed very efficiently in RFL-6 cells, both mutants expressed less efficiently (Supplementary Figure 2A). For this reason, rather than assessing MT stability biochemically at the population level (via Western blot, for example), we evaluated MT stability in cells that expressed comparable levels (as determined by anti-FLAG IF staining, Supplementary Figure 2B) of either UNC-45A or its mutants as determined by IF. Specifically, following overexpression of UNC-45A and its C-terminally or N-terminally deleted mutants, cells were fixed, extracted and stained with anti-alpha tubulin antibody 24 hours post infection. As shown in Figure 6A, we found that ectopic expression of both UNC-45A and its C-terminally deleted mutant resulted with significant decrease in total MT mass as compared to both control and N-terminally deleted cells. Quantification of MT mass per each condition is given in Figure 6B. We also looked at the possible effects of overexpression of UNC-45A and its mutant on MT breakages. While the numbers did not reach statistical significance (possibly due to the less efficient expression of the FLAG-tagged proteins used for these experiments and compared to the GFP-tagged proteins used in Figure 5) we found a clear trend of breakages being more frequent in WT- and dC-UNC-45A expressing cells as compared to control and dN cells (Figure 6C and D).

**Figure 6.**
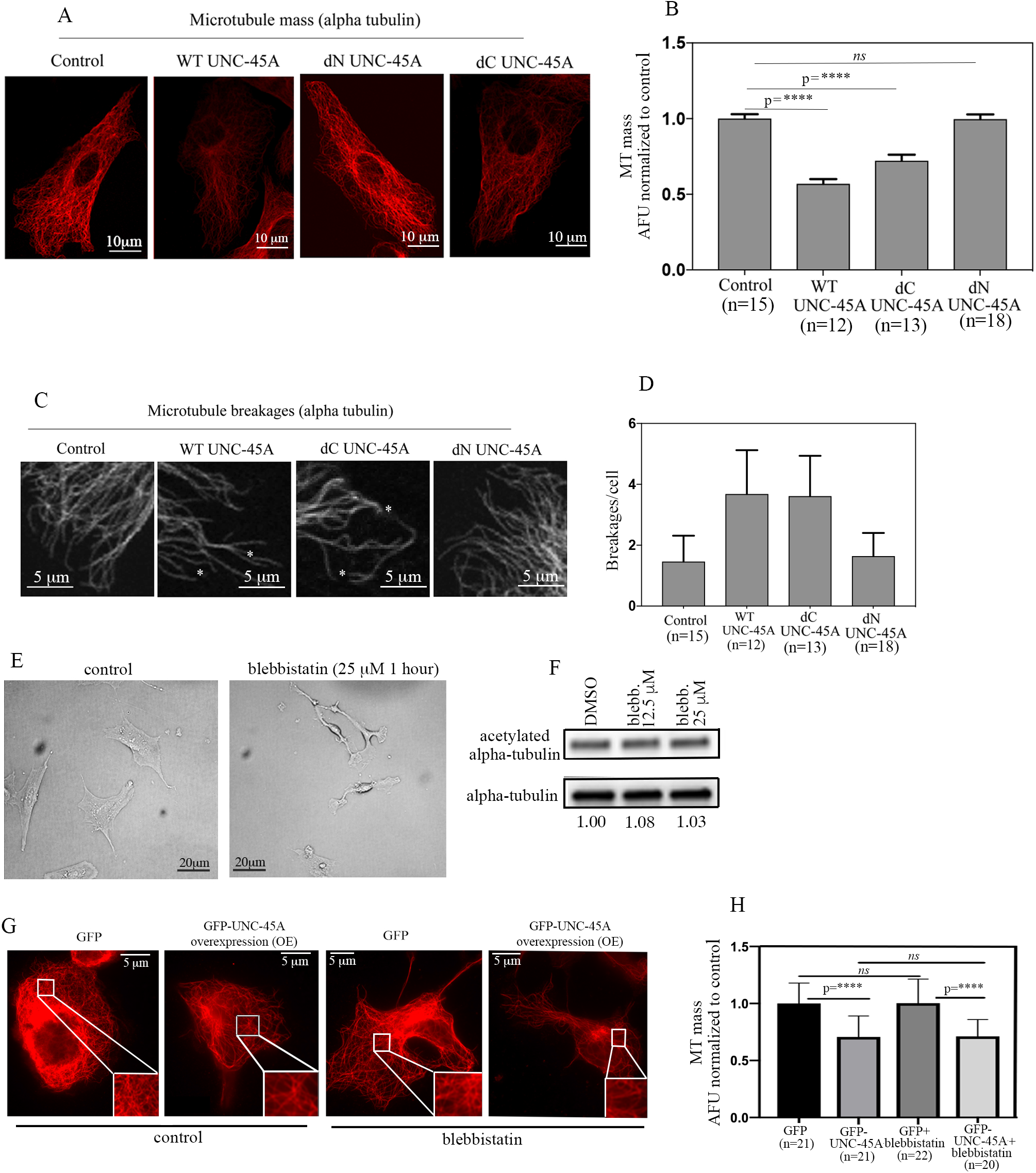
UNC-45A weakens and breaks MT in absence of its C-terminal NMII-Binding Domain and in presence of the NMII inhibitor blebbistatin. A. Representative images of RFL6 cells transduced with either empty vector (FLAG), Wild Type UNC-45A (WT), C-terminally deleted UNC-45A (dC), or N-terminally deleted UNC-45A (dN) FLAG-tagged proteins and stained for alpha tubulin (red). All images were taking using the same exposure time. B. Quantification of MT mass per each condition expressed as Arbitrary Fluorescence Units (AUF). n=number of cells analyzed per each condition (3 different areas per cell were evaluated). C. Representative images of RFL6 cells transduced with either empty vector (FLAG), Wild Type UNC-45A (WT), C-terminally deleted UNC-45A (dC), or N-terminally deleted UNC-45A (dN) FLAG tagged proteins stained for alpha tubulin (red) to visualize MTs and their breakages. All images were taken using the same exposure time. Black and white images are shown for better visualization. Asterisks indicate MT breakages. D. Quantification of MT breakages per cell per each condition. n=number of cells analyzed per each condition. Values nonsignificant. E. Representative image of RFL-6 cells either mock treated or treated with the indicated concentration of blebbistatin over an hour. F. Levels of acetylated tubulin versus total tubulin (numbers indicate the ratio) in control (DMSO) versus blebbistatin treated RFL-6 cells. G. Representative images of MT mass in control (GFP) and GFP-UNC-45A overexpressing RFL-6 cells (24 hours post infection) with or without blebbistatin treatment (25 μM for 1 hour) after staining with an anti-alpha-tubulin antibody to visualize MTs. All images were taking using the same exposure time. H. Quantification of MT mass per each condition. n= number of cells evaluated per each condition (3 different areas per cell were evaluated).

To directly confirm that in cells, UNC-45A destabilizes MTs independent of its effect on NMII, we investigated whether the effects of UNC-45A on MT destabilization (Figures 5 and 6) occurs even in the presence of the NMII inhibitor blebbistatin (Allingham et al., 2005). For these experiments, we first treated RFL-6 cells with either control DMSO or 12.5 and 25 μM of the NMII inhibitor blebbistatin over the period of 1 hour. The amount and duration of this treatment has been shown to potently and specifically inhibit NMII in a number of cell types (Kovacs et al., 2004). As expected and previously shown (Kovacs et al., 2004), blebbistatin treatment resulted with profound changes in cell morphology which is consistent with its effect on decreasing actomyosin contractility (Figure 6E). We then biochemically determined whether blebbistatin treatment lead to changes in the levels of acetylated alpha tubulin. We found no differences in the levels of acetylated alpha-tubulin in treated versus untreated cells (Figure 6F). This indicates that unlike what happens following UNC-45A KD (Figure 5A and (Mooneyham et al., 2019)), inhibition of NMII does not lead to an increase in the levels of the more stable MT population. Next, we reasoned that if UNC-45A destabilizes MTs independent of its effect on myosin then it should still have an effect when NMII is inhibited. To this end, GFP and GFP-UNC-45A expressing RFL-6 cells were treated with or without 25 μM of blebbistatin over a period of 1 hour. Cells were next fixed, extracted and stained with anti-alpha tubulin antibody 24 hours. As shown in Figure 6G, we found that ectopic expression of UNC-45A (as visualized by green channel, Supplementary Figure 2C), resulted with a significant decrease in total MT mass as compared to control even in presence of blebbistatin treatment. Quantification of MT mass per each condition is given in Figure 6H. Taken together this indicates that UNC-45A destabilizes MT independent of its effects on NMII activity.

## Discussion

MTs are highly dynamic components of the cytoskeleton and crucial for a variety of cellular functions. A number of MAPs are involved in regulating MT stability via either promoting MTs polymerization or depolymerization (Goodson and Jonasson, 2018). There are three main classes of MT destabilizing proteins: sequestering, plus-end depolymerizing and severing. MT sequestering proteins do not bind to MTs but cause their destabilization by binding to the pool of free tubulin in the cells which becomes then unavailable for MT polymerization (Belmont and Mitchison, 1996). MT plus-end depolymerizing proteins can either bind to the plus-end of the MT directly or bind along the length of MT and then travel to their plus-end, where they cause destabilization by either preventing incorporation of tubulin or by causing morphological changes that cause MT depolymerization for the plus-end (Howard and Hyman, 2007). MT severing proteins on the other hand, use ATP hydrolysis to “wedge” or “pull” tubulin dimers off the body of MTs (Roll-Mecak and McNally, 2010). This leads to a local destabilization of the MTs which is followed by breakages of the MTs along their lattice and creation of new plus ends from which further rapid depolymerization occurs. Katanin, the first MT severing protein discovered, owes its name to its ability to “cut” MTs along their length in the same fashion a sword (katana, in Japanese) would do (Quarmby, 2000).

We have previously shown that UNC-45A is a MAP with MT destabilizing properties *in vitro* and *in vivo* in a variety of mammalian cells and that it is able to counteract the MT stabilizing effects of taxol in cancer cells (Habicht et al., 2019a; Mooneyham et al., 2018). Here we show that *in vitro* UNC-45A binds to MTs lattice and causes bends and breakages along the length of paclitaxel-stabilized MTs. This is followed by MT depolymerization in a dosedependent manner. Furthermore, we show that UNC-45A overexpression causes a dramatic loss of MT mass and increase in MT breakages in RFL-6 cells. Importantly, RFL-6 cells lack the well-known MT stabilizing protein tau which has been recently proposed to prevent the activity of MT severing proteins (Baas and Qiang, 2019; Tan et al., 2019). Taken together our data suggests that UNC-45A may act via interacting with the MT lattice and causing its destabilization which leads to MT breakages and destabilization. Although additional studies are needed to understand the exact mechanism through which UNC-45A breaks MTs, it is likely that the mechanism is different than that of MT severing proteins. In fact, UNC-45A has no known ATP-ase domain and is capable of destabilizing paclitaxel-stabilized MTs *in vitro* independent of ATP (Bailey et al., 2016). Furthermore, and perhaps as a consequence of its ATP-independence, UNC-45A requires higher concentration and longer time as compared to other ATP-dependent MT destabilizing proteins to destabilize paclitaxel-stabilized MT *in vitro* (Jiang et al., 2017). This is also consistent with the fact that in cells, UNC-45A is approx. 20-fold more abundant (0.4 uM) than other MT destabilizing proteins, suggesting that UNC-45A may have a dominant role in the regulation of cellular MTs (Itzhak et al., 2016). Interestingly, MT depolymerizing proteins are known to exist as oligomers (including dimers, hexamers and dodecameric stacked rings), and their oligomerization is required for MT binding and/or MT depolymerization (Hertzer et al., 2006; Lupas and Martin, 2002; Neuwald et al., 1999; Sharp and Ross, 2012; Vale, 1991). Studies conducted using UNC-45 revealed that the protein exists as oligomers (dimers, tetramers, pentamers in *C.elegans* and dodecamers in *Drosophila) in vitro* and *in vivo(Gazda et al., 2013).* The sequence similarity between human and worm UNC-45A is over 50%, and the homology between human and fly UNC-45A is over 60%. This suggests that similarly to known MT severing proteins, UNC-45A may exist and act on MTs in an oligomeric state.

Interactions between actomyosin and MT systems are well known and many cytoskeletal-associated proteins are known to independently regulate both. The calcium binding family member SA10004A for instance, is overexpressed in a number of human cancers where it promotes cancer metastasis (Bresnick et al., 2015). This effect seems due to both, the fact that S1004A binds to NMIIA and promotes its depolymerization, and its myosin II-independent effect consistent with MT depolymerization (Dulyaninova et al., 2018). Furthermore, one of the most well-known cell cycle regulators, the cyclin-dependent kinase inhibitor 1B (p27), is both a MT- and a NMII-associated and co-localizing protein and has been show to regulate independently regulate both NMII activity and MTs stability in normal and cancer cells (Baldassarre et al., 2005; Besson et al., 2004; Bezanilla et al., 2000; Fabris et al., 2015; Godin et al., 2012; Murthy and Wadsworth, 2005; Serres et al., 2012). Similarly to, and perhaps because of its worm homolog UNC-45, UNC-45A has been mostly studied in the context of its role in regulating NMII activity. This includes studies from our and other groups showing that UNC-45A binds to and co-localizes with NMII and regulates its activity including the formation of stress fibers (Bazzaro et al., 2007; Guo et al., 2011; Iizuka et al., 2015; Iizuka et al., 2017a; Lehtimaki et al., 2017). Over the past five years however, it became evident that UNC-45A also regulates MT stability. This not only includes our above mentioned work (Habicht et al., 2019a; Mooneyham et al., 2018) but also work from others showing that UNC-45A co-localizes and cofractionates with gamma tubulin, the main component of the Microtubule Organizing Center (MTOC) (Jilani et al., 2015) and is abundant in subcellular fractions containing other MAP with MT destabilizing properties including Katanin and MCAK (Itzhak et al., 2016).

We had previously shown that UNC-45A binds MTs *in vitro* in the absence of NMII (Mooneyham et al., 2019). Here we show that in the same *in vitro* system UNC-45A binds to MTs even in the absence of its C-terminal NMII-binding domain. Further, we show that in a turbidimetric assay, UNC-45A lacking its C-terminal NMII-binding assay is still able to interfere with net MT polymerization in a time and dose-dependent fashion similarly to UNC-45A wildtype. Because in this particular assay tubulin polymerization occurs spontaneously and it is not promoted nor maintained by taxol (Shelanski et al., 1973), this result suggests that UNC-45A does not destabilizes MTs by simply competing or interfering with taxol binding to MTs. Also, interestingly, in this turbidimetric assay UNC-45A and its mutant lacking the C-terminal deleted myosin II do not interfere with the elongation phase of the MTs as nocodazole does, but, rather, they decrease the steady state levels of polymerized MTs. This is in accordance with our previous results (Mooneyham et al., 2019) indicating that UNC-45A does not destabilize MTs by sequestering free tubulin but it rather destabilizes already formed MTs. We also show that both, *in vitro* and in U2OS cells, UNC-45A lacking the C-terminal myosin II binding domain is still able to bind and destabilize interphase MTs similarly to UNC-45 wild-type. Importantly, this is the same cell type where UNC-45A was found to localize at the MTOC, to co-fractionate with gamma-tubulin, and to track with MTs close to the MTOC in metaphase cells (Jilani et al., 2015).

The effects of actomyosin system on MT stability are generally modest and in reverse direction. Specifically, while inhibition of myosin II leads to modest increase in MT stability (Kadir et al., 2011) we have previously shown that loss of UNC-45A leads to increase myosin II activity(Iizuka et al., 2017a). Here we show that treatment with the NMII inhibitor blebbistatin induces a profound change in the morphology of RLF-6 cells with cells becoming more elongated due to a reduction in actomyosin contractility (Allingham et al., 2005; Kovacs et al., 2004). This however, did not result in a change in the expression levels of acetylated alphatubulin suggesting that at least in this cell type, loss of NMII activity is not accompanied by increased MT stability. Lastly, we show that in RFL-6 cells UNC-45A destabilizes MTs even in presence of blebbistatin, confirming that UNC-45A acts on MTs independent of its effect/s on NMII and have a dual non-mutually exclusive role in regulating actomyosin and MT stability.

## Author contributions

Juri Habicht, Ashley Mooneyham, Asumi Hoshino, Mihir Shetty, Xiaonan Zhang, Edith Emmings, Qing Yang, and Courtney Coombes, performed the experiments and analyzed the data. Juri Habicht, Ashley Mooneyham, Melissa Gardner and Martina Bazzaro designed the experiments and analyzed the data. Juri Habicht, Ashley Mooneyham, Melissa Gardner and Martina Bazzaro wrote the manuscript.

## Acknowledgments

We thank Drs. Jaakko Lehtimäki and Pekka Lappalainen from the Institute of Biotechnology, University of Helsinki, Helsinki, Finland for the generous gift of the UNC-45A KO U2OS clones. We thank Dr. Valentino Clemente (University of Minnesota) for the help with image analysis. We thank Guillermo Marques (University of Minnesota Imaging Center) for assistance with image analysis. This work was supported by Department of Defense Ovarian Cancer Research Program Grant OC160377, the Minnesota Ovarian Cancer Alliance, the Randy Shaver Cancer Research Funds and the NIH grant NIGMS R01-GM130800 to Martina Bazzaro. Melissa Gardner was supported by NIH grant NIGMS R35-GM126974. The funders had no role in study design, data collection and analysis, decision to publish or preparation of the manuscript. The authors declare no conflict of interests.

## References

Ahmad, F. J., Yu, W., McNally, F. J., and Baas, P. W. (1999). An essential role for katanin in severing microtubules in the neuron. J Cell Biol 145, 305–315.

Allingham, J. S., Smith, R., and Rayment, I. (2005). The structural basis of blebbistatin inhibition and specificity for myosin II. Nat Struct Mol Biol 12, 378–379.

Baas, P. W., and Qiang, L. (2019). Tau: It’s Not What You Think. Trends Cell Biol 29, 452–461.

Baas, P. W., Rao, A. N., Matamoros, A. J., and Leo, L. (2016). Stability properties of neuronal microtubules. Cytoskeleton (Hoboken) 73, 442–460.

Baas, P. W., and Sudo, H. (2010). More microtubule severing proteins: more microtubules. Cell Cycle 9, 2273.

Bailey, M. E., Jiang, N., Dima, R. I., and Ross, J. L. (2016). Invited review: Microtubule severing enzymes couple atpase activity with tubulin GTPase spring loading. Biopolymers 105, 547–556.

Baldassarre, G., Belletti, B., Nicoloso, M. S., Schiappacassi, M., Vecchione, A., Spessotto, P., Morrione, A., Canzonieri, V., and Colombatti, A. (2005). p27(Kip1)-stathmin interaction influences sarcoma cell migration and invasion. Cancer cell 7, 51–63.

Barral, J. M., Bauer, C. C., Ortiz, I., and Epstein, H. F. (1998). Unc-45 mutations in Caenorhabditis elegans implicate a CRO1/She4p-like domain in myosin assembly. J Cell Biol 143, 1215–1225.

Barral, J. M., Hutagalung, A. H., Brinker, A., Hartl, F. U., and Epstein, H. F. (2002). Role of the myosin assembly protein UNC-45 as a molecular chaperone for myosin. Science 295, 669–671.

Bazzaro, M., Santillan, A., Lin, Z., Tang, T., Lee, M. K., Bristow, R. E., Shih Ie, M., and Roden, R. B. (2007). Myosin II co-chaperone general cell UNC-45 overexpression is associated with ovarian cancer, rapid proliferation, and motility. Am J Pathol 171, 1640–1649.

Belmont, L. D., and Mitchison, T. J. (1996). Identification of a protein that interacts with tubulin dimers and increases the catastrophe rate of microtubules. Cell 84, 623–631.

Bernick, E. P., Zhang, P. J., and Du, S. (2010). Knockdown and overexpression of Unc-45b result in defective myofibril organization in skeletal muscles of zebrafish embryos. BMC Cell Biol 11, 70.

Besson, A., Gurian-West, M., Schmidt, A., Hall, A., and Roberts, J. M. (2004). p27Kip1 modulates cell migration through the regulation of RhoA activation. Genes & development 18, 862–876.

Bezanilla, M., Wilson, J. M., and Pollard, T. D. (2000). Fission yeast myosin-II isoforms assemble into contractile rings at distinct times during mitosis. Current biology: CB 10, 397–400.

Bresnick, A. R., Weber, D. J., and Zimmer, D. B. (2015). S100 proteins in cancer. Nat Rev Cancer 15, 96–109.

Bujalowski, P. J., Nicholls, P., and Oberhauser, A. F. (2014). UNC-45B chaperone: the role of its domains in the interaction with the myosin motor domain. Biophys J 107, 654–661.

Di Paolo, G., Antonsson, B., Kassel, D., Riederer, B. M., and Grenningloh, G. (1997). Phosphorylation regulates the microtubule-destabilizing activity of stathmin and its interaction with tubulin. FEBS Lett 416, 149–152.

Dulyaninova, N. G., Ruiz, P. D., Gamble, M. J., Backer, J. M., and Bresnick, A. R. (2018). S100A4 regulates macrophage invasion by distinct myosin-dependent and myosin-independent mechanisms. Mol Biol Cell 29, 632–642.

Dunn, K. W., Kamocka, M. M., and McDonald, J. H. (2011). A practical guide to evaluating colocalization in biological microscopy. Am J Physiol Cell Physiol 300, C723–742.

Epstein, H. F., and Thomson, J. N. (1974). Temperature-sensitive mutation affecting myofilament assembly in Caenorhabditis elegans. Nature 250, 579–580.

Etard, C., Behra, M., Fischer, N., Hutcheson, D., Geisler, R., and Strahle, U. (2007). The UCS factor Steif/Unc-45b interacts with the heat shock protein Hsp90a during myofibrillogenesis. Dev Biol 308, 133–143.

Etheridge, L., Diiorio, P., and Sagerstrom, C. G. (2002). A zebrafish unc-45-related gene expressed during muscle development. Dev Dyn 224, 457–460.

Fabris, L., Berton, S., Pellizzari, I., Segatto, I., D’Andrea, S., Armenia, J., Bomben, R., Schiappacassi, M., Gattei, V., Philips, M. R., et al. (2015). p27kip1 controls H-Ras/MAPK activation and cell cycle entry via modulation of MT stability. Proceedings of the National Academy of Sciences of the United States of America 112, 13916–13921.

Gazda, L., Pokrzywa, W., Hellerschmied, D., Lowe, T., Forne, I., Mueller-Planitz, F., Hoppe, T., and Clausen, T. (2013). The myosin chaperone UNC-45 is organized in tandem modules to support myofilament formation in C. elegans. Cell 152, 183–195.

Geach, T. J., and Zimmerman, L. B. (2010). Paralysis and delayed Z-disc formation in the Xenopus tropicalis unc45b mutant dicky ticker. BMC Dev Biol 10, 75.

Godin, J. D., Thomas, N., Laguesse, S., Malinouskaya, L., Close, P., Malaise, O., Purnelle, A., Raineteau, O., Campbell, K., Fero, M., et al. (2012). p27(Kip1) is a microtubule-associated protein that promotes microtubule polymerization during neuron migration. Developmental cell 23, 729–744.

Goodson, H. V., and Jonasson, E. M. (2018). Microtubules and Microtubule-Associated Proteins. Cold Spring Harb Perspect Biol 10.

Guo, W., Chen, D., Fan, Z., and Epstein, H. F. (2011). Differential turnover of myosin chaperone UNC-45A isoforms increases in metastatic human breast cancer. J Mol Biol 412, 365–378.

Habicht, J., Mooneyham, A., Shetty, M., Zhang, X., Shridhar, V., Winterhoff, B., Zhang, Y., Cepela, J., Starr, T., Lou, E., and Bazzaro, M. (2019a). UNC-45A is preferentially expressed in epithelial cells and binds to and co-localizes with interphase MTs. Cancer Biol Ther, 1–10.

Habicht, J., Mooneyham, A., Shetty, M., Zhang, X., Shridhar, V., Winterhoff, B., Zhang, Y., Cepela, J., Starr, T., Lou, E., and Bazzaro, M. (2019b). UNC-45A is preferentially expressed in epithelial cells and binds to and co-localizes with interphase MTs. Cancer Biol Ther 20, 1304–1313.

Hartman, J. J., Mahr, J., McNally, K., Okawa, K., Iwamatsu, A., Thomas, S., Cheesman, S., Heuser, J., Vale, R. D., and McNally, F. J. (1998). Katanin, a microtubule-severing protein, is a novel AAA ATPase that targets to the centrosome using a WD40-containing subunit. Cell 93, 277–287.

Hartman, J. J., and Vale, R. D. (1999). Microtubule disassembly by ATP-dependent oligomerization of the AAA enzyme katanin. Science 286, 782–785.

Hertzer, K. M., Ems-McClung, S. C., Kline-Smith, S. L., Lipkin, T. G., Gilbert, S. P., and Walczak, C. E. (2006). Full-length dimeric MCAK is a more efficient microtubule depolymerase than minimal domain monomeric MCAK. Mol Biol Cell 17, 700–710.

Howard, J., and Hyman, A. A. (2007). Microtubule polymerases and depolymerases. Curr Opin Cell Biol 19, 31–35.

Hutagalung, A. H., Landsverk, M. L., Price, M. G., and Epstein, H. F. (2002). The UCS family of myosin chaperones. J Cell Sci 115, 3983–3990.

Iizuka, Y., Cichocki, F., Sieben, A., Sforza, F., Karim, R., Coughlin, K., Isaksson Vogel, R., Gavioli, R., McCullar, V., Lenvik, T., et al. (2015). UNC-45A Is a Nonmuscle Myosin IIA Chaperone Required for NK Cell Cytotoxicity via Control of Lytic Granule Secretion. Journal of immunology 195, 4760–4770.

Iizuka, Y., Mooneyham, A., Sieben, A., Chen, K., Maile, M., Hellweg, R., Schutz, F., Teckle, K., Starr, T., Thayanithy, V., et al. (2017a). UNC-45A is required for neurite extension via controlling NMII activation. Mol Biol Cell 28, 1337–1346.

Iizuka, Y., Mooneyham, A., Sieben, A., Chen, K., Maile, M., Hellweg, R., Schutz, F., Teckle, K., Starr, T., Thayanithy, V., et al. (2017b). UNC-45A is Required for Neurite Extension via Controlling NMII activation. Molecular biology of the cell.

Itzhak, D. N., Tyanova, S., Cox, J., and Borner, G. H. (2016). Global, quantitative and dynamic mapping of protein subcellular localization. Elife 5.

Jiang, K., Rezabkova, L., Hua, S., Liu, Q., Capitani, G., Altelaar, A. F. M., Heck, A. J. R., Kammerer, R. A., Steinmetz, M. O., and Akhmanova, A. (2017). Microtubule minus-end regulation at spindle poles by an ASPM-katanin complex. Nat Cell Biol 19, 480–492.

Jilani, Y., Lu, S., Lei, H., Karnitz, L. M., and Chadli, A. (2015). UNC45A localizes to centrosomes and regulates cancer cell proliferation through ChK1 activation. Cancer Lett 357, 114–120.

Joo, E. E., and Yamada, K. M. (2014). MYPT1 regulates contractility and microtubule acetylation to modulate integrin adhesions and matrix assembly. Nat Commun 5, 3510.

Kadir, S., Astin, J. W., Tahtamouni, L., Martin, P., and Nobes, C. D. (2011). Microtubule remodelling is required for the front-rear polarity switch during contact inhibition of locomotion. J Cell Sci 124, 2642–2653.

Kovacs, M., Toth, J., Hetenyi, C., Malnasi-Csizmadia, A., and Sellers, J. R. (2004). Mechanism of blebbistatin inhibition of myosin II. J Biol Chem 279, 35557–35563.

Lee, C. F., Melkani, G. C., Yu, Q., Suggs, J. A., Kronert, W. A., Suzuki, Y., Hipolito, L., Price, M. G., Epstein, H. F., and Bernstein, S. I. (2011). Drosophila UNC-45 accumulates in embryonic blastoderm and in muscles, and is essential for muscle myosin stability. J Cell Sci 124, 699–705.

Lehtimaki, J. I., Fenix, A. M., Kotila, T. M., Balistreri, G., Paavolainen, L., Varjosalo, M., Burnette, D. T., and Lappalainen, P. (2017). UNC-45a promotes myosin folding and stress fiber assembly. J Cell Biol 216, 4053–4072.

Lou, E., Zhai, E., Sarkari, A., Desir, S., Wong, P., Iizuka, Y., Yang, J., Subramanian, S., McCarthy, J., Bazzaro, M., and Steer, C. J. (2018). Cellular and Molecular Networking Within the Ecosystem of Cancer Cell Communication via Tunneling Nanotubes. Front Cell Dev Biol 6, 95.

Lupas, A. N., and Martin, J. (2002). AAA proteins. Current opinion in structural biology 12, 746–753.

Mooneyham, A., Iizuka, Y., Yang, Q., Coombes, C., McClellan, M., Shridhar, V., Emmings, E., Shetty, M., Chen, L., Ai, T., et al. (2018). UNC-45A Is a Novel Microtubule-Associated Protein and Regulator of Paclitaxel Sensitivity in Ovarian Cancer Cells. Mol Cancer Res.

Mooneyham, A., Iizuka, Y., Yang, Q., Coombes, C., McClellan, M., Shridhar, V., Emmings, E., Shetty, M., Chen, L., Ai, T., et al. (2019). UNC-45A Is a Novel Microtubule-Associated Protein and Regulator of Paclitaxel Sensitivity in Ovarian Cancer Cells. Mol Cancer Res 17, 370–383.

Murthy, K., and Wadsworth, P. (2005). Myosin-II-dependent localization and dynamics of Factin during cytokinesis. Current biology: CB 15, 724–731.

Neuwald, A. F., Aravind, L., Spouge, J. L., and Koonin, E. V. (1999). AAA+: A class of chaperone-like ATPases associated with the assembly, operation, and disassembly of protein complexes. Genome research 9, 27–43.

Ng, D. C., Lin, B. H., Lim, C. P., Huang, G., Zhang, T., Poli, V., and Cao, X. (2006). Stat3 regulates microtubules by antagonizing the depolymerization activity of stathmin. J Cell Biol 172, 245–257.

Ni, W., Hutagalung, A. H., Li, S., and Epstein, H. F. (2011). The myosin-binding UCS domain but not the Hsp90-binding TPR domain of the UNC-45 chaperone is essential for function in Caenorhabditis elegans. J Cell Sci 124, 3164–3173.

Qiang, L., Yu, W., Andreadis, A., Luo, M., and Baas, P. W. (2006). Tau protects microtubules in the axon from severing by katanin. J Neurosci 26, 3120–3129.

Quarmby, L. (2000). Cellular Samurai: katanin and the severing of microtubules. J Cell Sci 113 (Pt 16), 2821–2827.

Ritter, B., Ferguson, S. M., De Camilli, P., and McPherson, P. S. (2017). A lentiviral system for efficient knockdown of proteins in neuronal cultures [version 1; referees: 2 approved]. MNI Open Res 1.

Roll-Mecak, A., and McNally, F. J. (2010). Microtubule-severing enzymes. Curr Opin Cell Biol 22, 96–103.

Serres, M. P., Kossatz, U., Chi, Y., Roberts, J. M., Malek, N. P., and Besson, A. (2012). p27(Kip1) controls cytokinesis via the regulation of citron kinase activation. The Journal of clinical investigation 122, 844–858.

Sharp, D. J., and Ross, J. L. (2012). Microtubule-severing enzymes at the cutting edge. Journal of cell science 125, 2561–2569.

Shelanski, M. L. (1973). Chemistry of the filaments and tubules of brain. J Histochem Cytochem 21, 529–539.

Shelanski, M. L., Gaskin, F., and Cantor, C. R. (1973). Microtubule assembly in the absence of added nucleotides. Proc Natl Acad Sci U S A 70, 765–768.

Shi, H., and Blobel, G. (2010). UNC-45/CRO1/She4p (UCS) protein forms elongated dimer and joins two myosin heads near their actin binding region. Proc Natl Acad Sci U S A 107, 21382–21387.

Sudo, H., and Baas, P. W. (2010). Acetylation of microtubules influences their sensitivity to severing by katanin in neurons and fibroblasts. J Neurosci 30, 7215–7226.

Sudo, H., and Baas, P. W. (2011). Strategies for diminishing katanin-based loss of microtubules in tauopathic neurodegenerative diseases. Hum Mol Genet 20, 763–778.

Tan, R., Lam, A. J., Tan, T., Han, J., Nowakowski, D. W., Vershinin, M., Simo, S., Ori-McKenney, K. M., and McKenney, R. J. (2019). Microtubules gate tau condensation to spatially regulate microtubule functions. Nat Cell Biol 21, 1078–1085.

Vale, R. D. (1991). Severing of stable microtubules by a mitotically activated protein in Xenopus egg extracts. Cell 64, 827–839.

