## Supplementary figures and images for "UNC-45A Weakens and Breaks MT Lattice Independent of its Effect on Non-Muscle Myosin II"

### Supplementary Figure 1

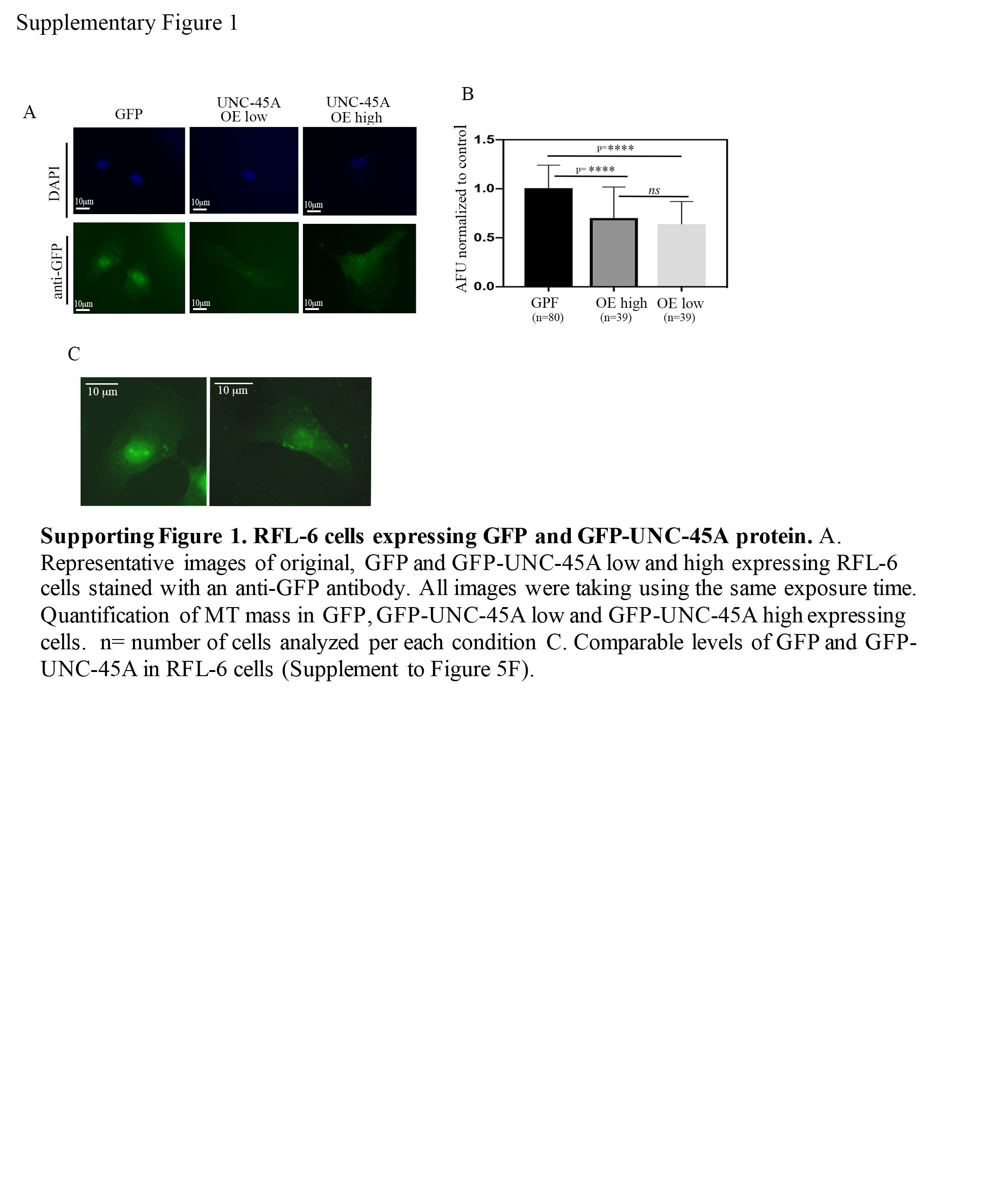

### Supplementary Figure 2

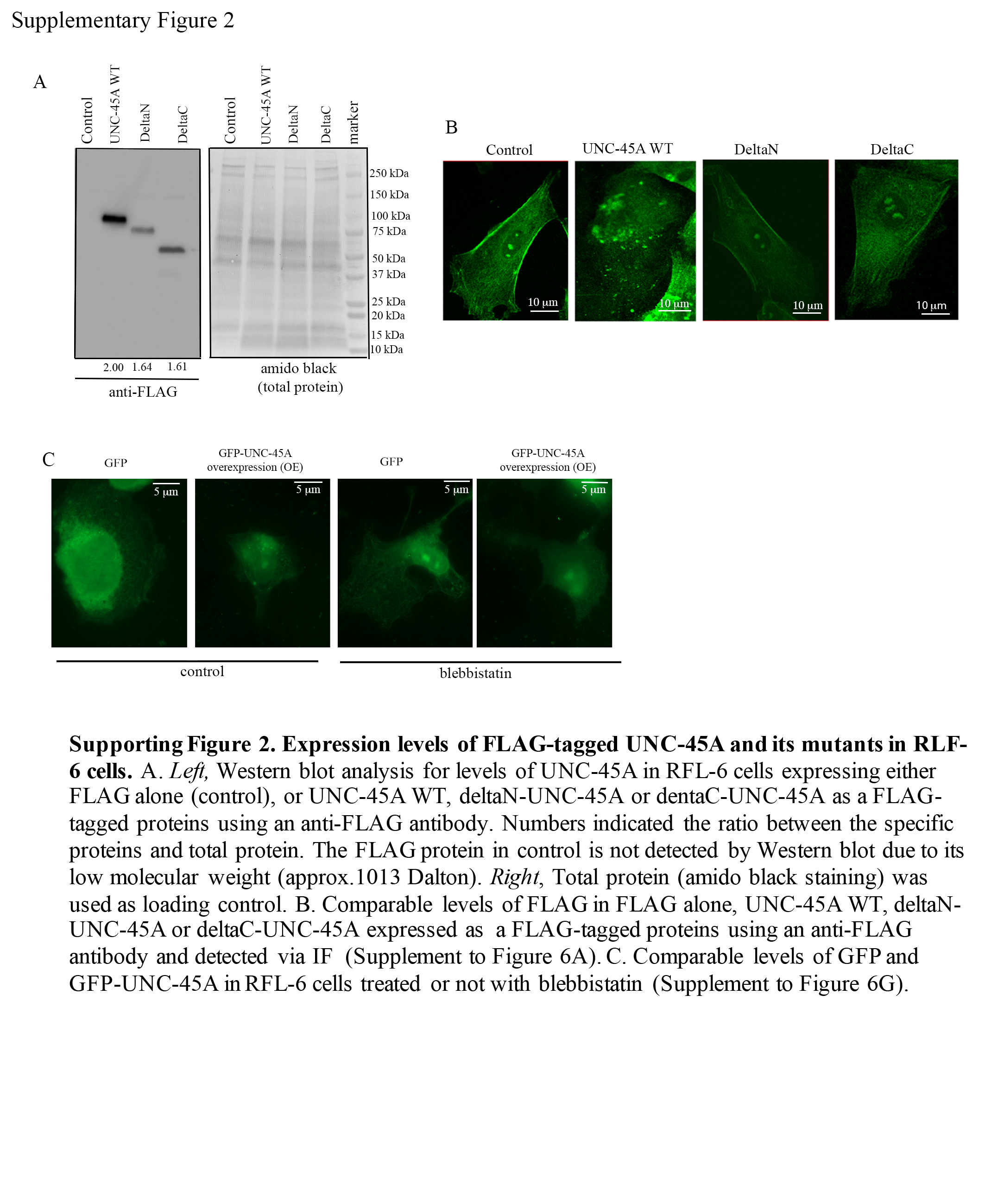
